# Characterization of Diverse Homoserine Lactone Synthases in *Escherichia coli*

**DOI:** 10.1101/279349

**Authors:** René Daer, Cassandra Barrett, Ernesto Luna Melendez, Jiaqi Wu, Stefan Tekel, Jimmy Xu, Brady Dennison, Ryan Muller, Karmella Haynes

**Affiliations:** Arizona State University, School of Biological and Health Systems Engineering, 501 E. Tyler Mall, ECG 344A, Tempe, AZ, USA; School of Molecular Sciences, Arizona State University, Tempe, AZ; School of Computing, Informatics, and Decision Systems Engineering, Arizona State University, Tempe, AZ; School of Life Sciences, Arizona State University, Tempe, AZ; School of Engineering of Matter, Transport, and Energy, Arizona State University, Tempe, AZ; Department of Chemistry and Biochemistry, Arizona State University, Tempe, AZ, USA

**Keywords:** quorum sensing, homoserine lactone, functional orthogonality, HSL synthase

## Abstract

Quorum sensing networks have been identified in over one hundred bacterial species to date. A subset of these networks regulate group behaviors, such as bioluminescence, virulence, and biofilm formation, by sending and receiving small molecules called homoserine lactones (HSLs). Bioengineers have incorporated quorum sensing pathways into genetic circuits to connect logical operations. However, the development of higher-order genetic circuitry is inhibited by crosstalk, in which one quorum sensing network responds to HSLs produced by a different network. Here, we report the construction and characterization of a library of ten synthases including some that are expected to produce HSLs that are incompatible with the Lux pathway, and therefore show no crosstalk. We demonstrated their function in a common lab chassis, *Escherichia coli* BL21, and in two contexts, liquid and solid agar cultures, using decoupled Sender and Receiver pathways. We observed weak or strong stimulation of a Lux Receiver by longer-chain or shorter-chain HSL-generating Senders, respectively. We also considered the under-investigated risk of unintentional release of incompletely deactivated HSLs in biological waste. We found that HSL-enriched media treated with bleach is still bioactive, while autoclaving deactivates LuxR induction. This work represents the most extensive comparison of quorum sensing synthases to date and greatly expands the bacterial signaling toolkit while recommending practices for disposal based on empirical, quantitative evidence.

## Introduction

Quorum sensing networks enable bacteria to monitor and respond to changes in their population density by coupling gene regulation with diffusible chemical signals from neighboring bacteria [1]. These signaling networks control group behaviors such as virulence, biofilm formation, and motility [2]. One class of these chemical signals, known as homoserine lactones (HSLs), is produced by a family of synthase enzymes called LuxI-like proteins [3–5]. HSLs have traditionally been referred to as N-acyl homoserine lactones (AHLs). However synthases also produce non-acyl chain homoserine lactones, so in this report we use the general term HSL. Accumulation of HSLs results in activation of a DNA binding, regulator protein, or LuxR-like protein, that controls expression of genes (Fig 1A) involved in microbial population behavior. Homologous HSL networks have been identified in over one hundred species of bacteria and the discovery of more networks is ongoing [6]. Each network includes an HSL synthase protein that catalyzes the synthesis of specific HSL signaling molecules (Fig 1B), a regulator that is allosterically regulated by the HSL ligand, and a promoter that typically contains a palindromic sequence that is bound by the HSL-regulator complex.

**Figure 1:**
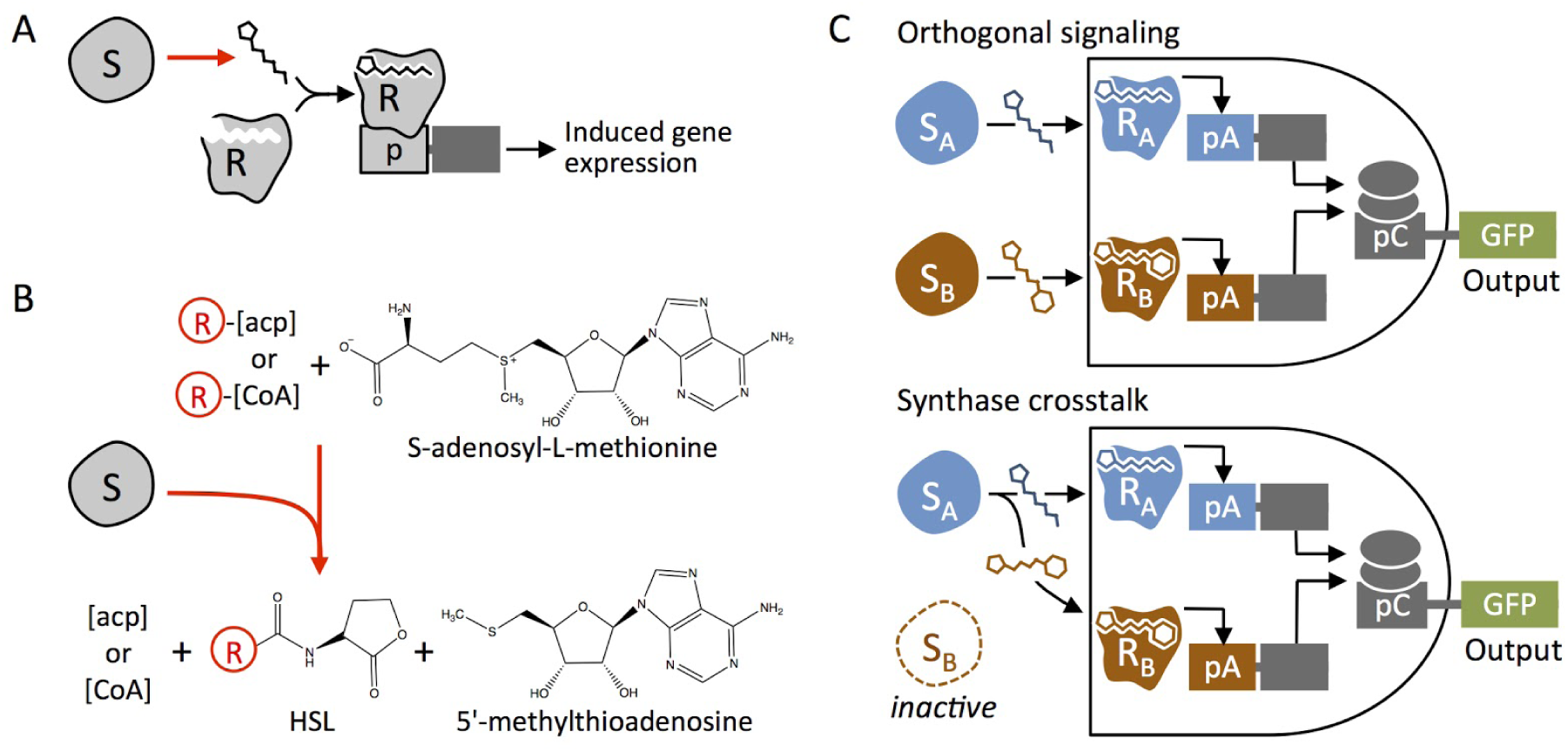
Homoserine lactone (HSL) function, production, and role in quorum sensing crosstalk. (A) Synthase protein (S) catalyzes formation of the HSL species. Regulator protein (R) complexes with the HSL ligand and binds to the promoter, which induces expression of downstream genes. (b) Synthase proteins catalyze the production of HSL molecules by facilitating lactone ring formation from S-adenosyl methionine and attaching a variable R-group tail to the lactone ring from one of two donors, acyl-carrier proteins (ACP) or coenzyme A (CoA). (c) The cartoon illustrates an example of the consequence of crosstalk. Two pathways, SA > RA and SB > RB are designed to operate in parallel as an AND gate sensor. When each synthase generates a single predominant HSL, the output indicates the activity of both S_A_ and S_B_ as intended. However, if S_A_ produces high levels of primary and secondary products that stimulate both RA and RB, a fault occurs where output is produced although the AND condition is not met.

Scientists have taken advantage of the simplicity of these systems to incorporate signal processing pathways into gene circuits as genetic wires to convert an output from one computation into an input of another. In an engineered system, the synthase protein can be considered a “Sender” module which produces the input for a “Receiver module” comprised of the regulator and the inducible promoter upstream of an output, such as GFP [7]. Engineered quorum sensing networks are used for a variety of applications including metabolic engineering, computational circuits, and medicine. Engineered quorum sensing systems that incorporate HSL senders, rather than exogenously added synthetic HSLs, allow researchers to increase the computational complexity of a circuit. If a system employs multiple, non-overlapping quorum sensing networks, simultaneous parallel computation [7] can occur within a single cell or linked across populations in co-culture or in solid agar. Several examples of successful implementation of multiple networks in one circuit have been reported [8–13], however, these examples are limited to two networks in parallel and took significant efforts to optimize.

Promiscuity and crosstalk can occur at different steps in quorum sensing pathways, including non-specific interactions between the HSL-ligands and regulators, as well as interactions of Regulator-HSL complexes with promoters [14–17]. To mitigate this challenge, much effort has has been invested in identifying or engineering orthogonal quorum sensing networks [18]. In this study we address crosstalk due to context-specific synthase activity, where a single synthase can generate and unexpected profile of HSL molecules (Fig 1C).

Natural diversity arises from variability in the HSL signal or signals employed by each network, specifically, variability in the R-groups on the lactone rings [19]. Recent efforts have led to the identification and engineering of regulators with greater specificity for chemically-distinct HSL ligands [18]. Reliable implementation of these new tools without ligand crosstalk requires complementary synthases to generate the expected HSL and minimal or no secondary products. While many synthases are known to produce a single HSL product in their native species, it cannot be assumed that they will behave similarly when expressed exogenously. Transgenic HSL synthases use endogenous acyl-carrier protein (ACP) or coenzyme A (CoA) donors as substrates to catalyze HSL formation (Fig 1B). This suggests that even with proper protein expression and folding, production of the expected HSLs depends on the availability of the appropriate ACP or CoA donor, which varies across bacterial species [20–22]. Indeed, previous work has demonstrated that some HSL synthases produce multiple HSLs [7] and that the HSL production profile can differ depending on the chassis the synthase is expressed in [23]. It is important to test sender and receiver devices in the chassis and context relevant to the intended application.

Crosstalk between species of quorum sensing bacteria is prevalent in nature as well. Environmental bacteria can respond to HSLs produced by other species for both cooperative and competitive gains [24–26]. The generation and disposal of large quantities of HSLs could cause misregulation of quorum sensing within microbial niches. Quorum sensing networks are ubiquitous and crucial to many natural systems; they coordinate virulent group behaviors in human and other pathogens [24,27], maintain the exchange of nutrients between nitrogen-fixing Rhizobia and legumes [28], and regulate signalling between photosynthetic symbionts and their reef coral hosts [29]. It is therefore necessary to critically evaluate conventional methods for bacterial culture disposal to determine whether they are sufficient for deactivating HSLs in order to mitigate potential health risks and environmental impacts.

## Results and Discussion

### Construction and expression of a HSL synthase library

Minimizing crosstalk between quorum sensing networks will enable genetic engineers to operate quorum sensing pathways in parallel, and therefore build more complex circuits. Toward this end, we selected ten HSL synthase genes that are known produce chemically-diverse signaling molecules with few secondary products in their native species. We considered the length of the acyl chain, the chain saturation, and the number of different HSLs produced by a single synthase. From our previously published list of reported synthase proteins and their HSL profiles [30], we selected the following sythases: RpaI, BraI, RhlI, BjaI, EsaI, LuxI, AubI, LasI, and CerI (Table 1). This set represents HSLs with chain lengths from 3 to 18 carbons and modifications including phenol, phenyl, carbonyl, and methyl groups. We included LuxI as it is the most commonly used network and the cognate Sender to our LuxR receiver device. While the inclusion of EsaI may seem redundant as they both produce N-(3-oxo-hexanoyl)-l-homoserine lactone (3-oxo-C6-HSL) as their major HSL, Gould *et al.* showed that EsaI only produces 3-oxo-C6-HSL in *E. coli* and therefore may have no secondary HSL products [31]. We also included SinI despite its promiscuity because of the unique HSLs it synthesizes, including the unsaturated 3-oxo-7,8-cis-C16-HSL and the long chain N-octadecanoyl-l-homoserine lactone (C18-HSL) [32].

**Table 1:**
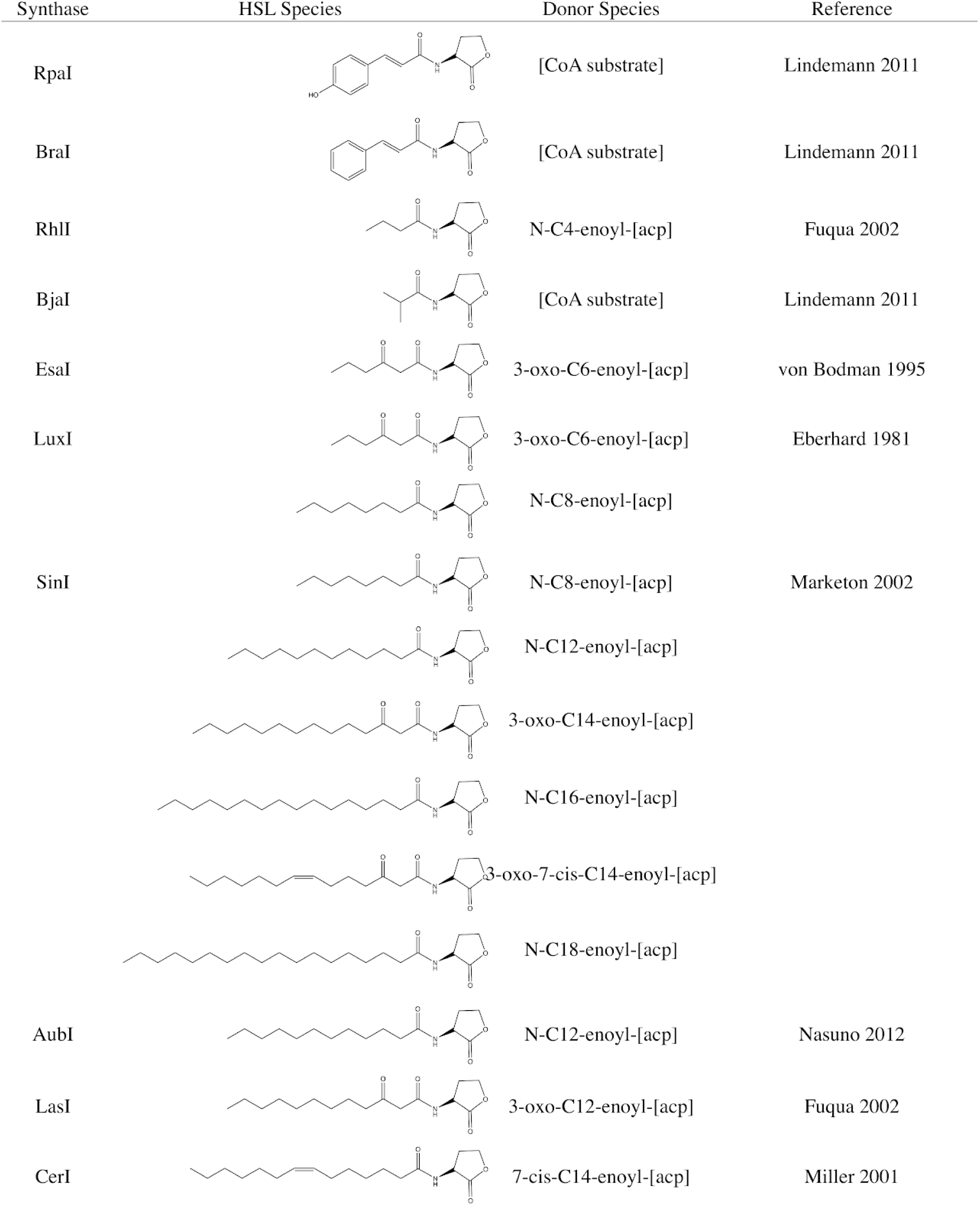
Synthases used in this study. The table lists the synthase symbol, the structures of the homoserine lactone (HSL) species produced by the native organism, and the acyl-carrier protein (ACP) or coenzyme A (CoA) donor species catalyzed by the synthase. [23,32–36]

We obtained synthesized DNA based on the publicly available sequences for each synthase open reading frame (ORF) (Supplemental Information). We designed and constructed a plasmid for simple directional cloning and high expression of HSL synthase homologues in *E. coli* (Fig 2A). The Sender vector (Bba_K2033011) includes a strong constitutive promoter (pTetR), a strong ribosome binding site (B0034), and high-copy number origin of replication (pUC19-derived pMB1, 100-300 copies per cell) for maximal protein expression and HSL production, with the option to switch to an inducible expression approach (in a chassis that expresses the TetR protein) [37]. The vector also carries a fluorescent protein ORF, mCherry, for pTetR-driven bicistronic expression such that mCherry signal indicates the production of full-length Synthase-mCherry mRNA (Fig 2A).

**Figure 2:**
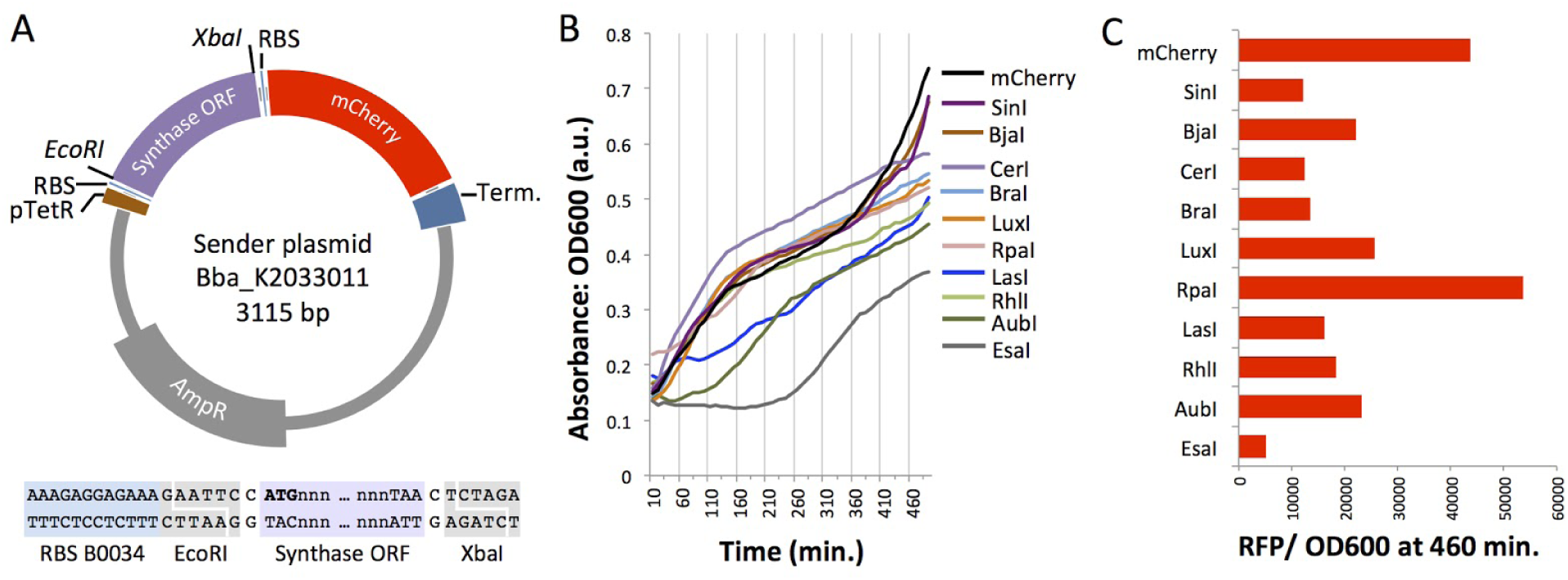
Sender plasmids expressed in *Escherichia coli* BL21. (A) The modular Sender vector allows facile cloning of any *EcoRI, XbaI*-flanked synthase open reading frame (ORF) and co-expression with mCherry. (B) Optical density (OD) readings for *E. coli* cultures expressing each of the Sender plasmids. (C) mCherry expression indicated by RFP signal normalized to OD600 over the 8 hour growth period.

We carried out fluorescence plate reader assays to monitor mCherry expression levels in BL21 E. coli that were transformed with one of each Sender variant. An mCherry-only plasmid (Bba_K2033011 without Synthase) was used as a positive control. While most of the Synthase-expressing cultures showed growth that tracked closely with the mCherry control, the lag in growth up to 110 or 260 minutes for EsaI, AubI, and LasI suggests metabolic burden or toxicity in these three cultures (Fig 2B). Red fluorescence protein (RFP) signal values normalized by absorbance (OD600) indicated that mCherry expression was roughly 12% to 50% that of the control, with the exception of RpaI. Incomplete transcription or less efficient expression from the larger Bba_K2033011 + Synthase plasmids (585 - 714 additional bp) may account for lower mCherry signals. An unidentified cryptic promoter within the RpaI ORF could have enhanced mCherry expression. Overall, these results validate the production of mRNA transcripts containing both the HSL synthase and the mCherry ORF and show that mCherry expression increases over time. We submitted to the iGEM Registry of Standard Parts sequences that were not previously represented in this public collection (Supplemental Table S1).

### Induction of a LuxR Receiver device with synthases from the Sender library

In order to measure Sender functionality, we induced a LuxR Receiver device with HSL-enriched, cell-free media from Sender cultures. The promiscuity of Lux signal-sensing is well established [19] and it is often used to test for the general presence of HSLs [11]. The BioBrick plasmid BBa_F2620 constitutively expresses the LuxR regulator protein and contains the inducible pLux promoter [19]. We cloned EGFP (BBa_E0240) downstream of pLux so that upon addition of a strong ligand (HSL), LuxR would bind the pLux promoter and induce EGFP expression. We transformed a common lab chassis, *E. coli* BL21 with our LuxR Receiver device, F2620_EGFP, or with each of the Sender plasmids. We grew the Senders and Receivers in separate liquid cultures, collected and filtered HSL-enriched media from Sender cultures, or mock-enriched media from mCherry synthase-minus bacteria, and treated Receiver cells with the filtered media (Fig 3A, Table 2). GFP signal and OD600 were measured over time (4 hours) for each sample using a plate reader.

**Table 2:** Preparation of Receiver induction experiments. The total volume of mixed media (Total, 150 μL) was added to 150 μL of Receiver liquid culture for a final volume of 300 μL.

| Sample | Sender media (μL) | Mock-enriched media (μL) | Total | Final |
| --- | --- | --- | --- | --- |
| Receiver + mCherry media | 0 | 150 | 150 | 300 |
| Receiver + 10% Sender media | 30 | 120 | 150 | 300 |
| Receiver + 25% Sender media | 75 | 75 | 150 | 300 |
| Receiver + 50% Sender media | 150 | 0 | 150 | 300 |

**Figure 3:**
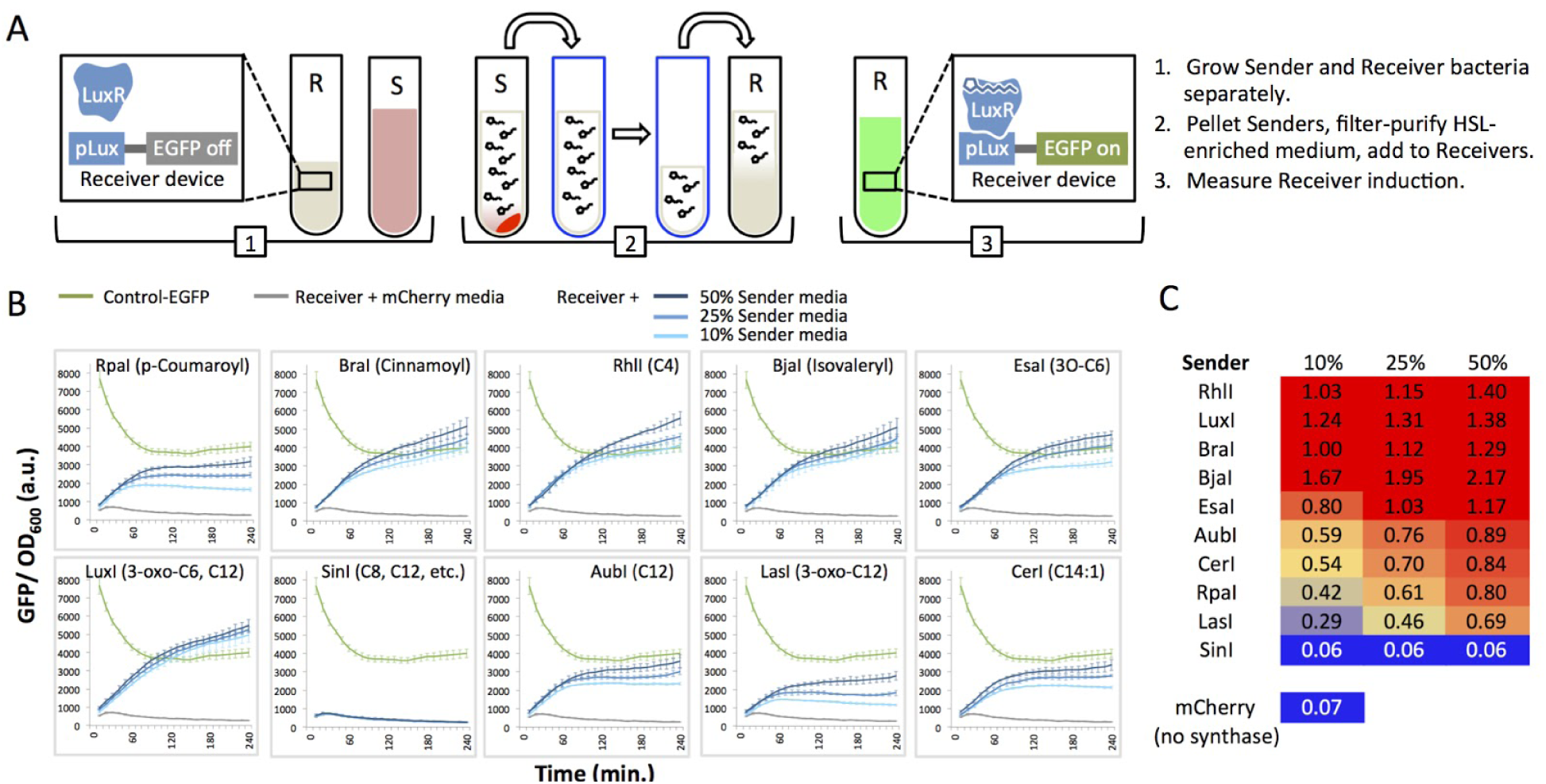
Induction of the LuxR Receiver with HSL-enriched media from Senders. (A) The cartoon illustrates the experimental approach for Receiver inductions. Sender (S) culture media is separated from pelleted Sender cells (arrow 1), filtered (arrow 2), and diluted (see Table 2), and used to induce Receiver cells. (B) GFP/OD600 of Sender-media-treated LuxR Receiver cells. EGFP and optical density (OD600) are measured every 10 minutes for 240 minutes (4 hours). Graphs show means of triplicate wells (bars, standard deviation). (C) The heat map shows GFP/ OD600 values after 240 min of induction, normalized to the GFP/ OD600 value for Control-GFP.

We considered results reported by Canton et al [38] as a model to predict and interpret the results from our Synthase-driven induction experiments. Their experiment established the range of responses and relative sensitivity of BBa_F2620 (LuxR) to known concentrations of homogenous HSL solutions. If we assume equal concentrations of homogenous HSL production from each *E. coli*-expressed synthase, we can hypothesize that the media will show relatively strong to weak (or no) stimulation of pLux-EGFP in the following order: LuxI (3-oxo-C6, C12) or EsaI (3-oxo-C6), SinI (C8, C12, and others), AubI (C12), RhlI (C4). We cannot make a prediction for the synthases that are not represented in the Canton et al experiment: RpaI (p-Coumaroyl), BraI (Cinnamoyl), BjaI (Isovaleryl), CerI (C14:1), and LasI (3O-C12). Our prediction is confounded by the possibility that transgenically expressed non-native synthases generate a mixture of active and inactive products at varying concentrations, which can be difficult to analyze quantitatively. Therefore, pLux-EGFP induction using synthetic HSLs has limited applicability for cell-based inducers.

We observed EGFP signal above background (Receivers plus mock-enriched media) for all but one of our Senders, *SinI* (Fig 3B). We can conclude from these data that nine synthases expressed bioactive, cell membrane permeable HSLs in *E. coli* BL21. We also measured signal from a constitutively-expressed pLac-EGFP (called “Control-EGFP” here) to establish a threshold for full activation. Control-EGFP cells showed high initial signal (GFP/OD600) that decreased over time and reached steady state after ∼90 min. At 240 minutes (6 hours) post-induction, Receivers treated with any of the three concentrations of Sender media (10%, 25%, 50%) from RhlI, LuxI, BraI, and BjaI generated GFP signal that reached Control-EGFP levels (Fig 3C, values ≥ 1.0). For EsaI, 25% and 50% Sender media was sufficient to induce full activation of EGFP. EsaI and LuxI both produce 3-oxo-C6-HSL in their native organisms [23,35] and *E. coli* [31] which strongly induces LuxR [38]. We expect RhlI to produce mostly N-butyryl-l-homoserine lactone (C4-HSL) which Canton *et al.* found to only induce this LuxR receiver at high concentrations [38]. However, Ortori *et al.* found that *Pseudomonas aeruginosa*, when mutated to only express RhlI, produced small amounts of N-hexanoyl-l-homoserine lactone (C6-HSL) and 3-oxo-C6-HSL [39]. The degree of LuxR response to RhlI that we observed is more consistent with this HSL profile and suggests that in *E. coli* BL21, RhlI produces C6-HSL and/or 3-oxo-C6-HSL. This result demonstrates the importance of testing synthases in context.

LasI, AubI, CerI, and RpaI showed induction above background at all dilutions, however, pLux-EGFP did not become fully activated (Fig 3B). These synthases are expected to produce HSLs with longer chains (Table 1). LuxR could interact poorly with these HSL ligands or the synthases could have low activity in our chassis.

### Characterization of sender induction in solid agar cultures

We next determined the behavior of our sender devices in a different context: spatially separated from the receiver device in solid agar cultures. We plated each sender in the center of a 10 cm agar plate and applied Receiver bacteria across an area spanning up to 3.0 cm from the Sender spot (Fig 4). After 16 hours of growth we measured the induction distance, the length from the center where the sender was plated to the edge of GFP expression by the LuxR receiver (Fig 4). The strongest inducers of LuxR in liquid culture, RhlI, LuxI, BraI, BjaI, and EsaI (Fig 3B), all induced LuxR on the agar plate. RpaI and CerI pLux-EGFP induction values were lower than those for the five strongest inducers in both experiments. Also consistent with the liquid induction experiment, SinI showed no induction of LuxR on the agar plate. In certain cases however, the relative performances of synthases were inconsistent in liquid versus agar cultures. For instance, EsaI induced pLux-EGFP over the longest distance overall, and almost two-fold the distance for the cognate synthase LuxI. However, EsaI did not induce the Receiver as strongly as LuxI in the liquid culture experiments. While LuxR responded to to LasI HSL-enriched liquid media in a dose-dependent manner (Fig 3B), we detected no EGFP signal over background on the agar plate (Fig 5B). Finally, the maximum value of AubI-induced pLux-EGFP (50% enriched media) was roughly half that of the strongest inducers (Fig 3C), whereas the distance of AubI-induced EGFP signal in the agar experiment (0.58 cm) fell within the range of values for the same inducers (BjaI, 0.41; RhlI, 0.66 cm) (Fig 4). Inconsistencies between the liquid and agar culture experiments might be due to HSL-specific diffusion limits in agar where there is no mechanical mixing, or differences in the stability of the HSLs in each medium. Furthermore, the agar experiment exposed Receiver bacteria to continual expression of each synthase from a growing population of Sender cells. In contrast, the fixed amount of available HSLs in the liquid culture experiments would be more affected by decay. Possible differences in enzyme-specific HSL production rates over the 16-hour period of plate incubation might account for the observed difference between LuxI and EsaI, which are expected to produce the same HSL. Overall, our results underscore the importance of testing devices in the chassis and context relevant to the application of interest.

**Figure 4:**
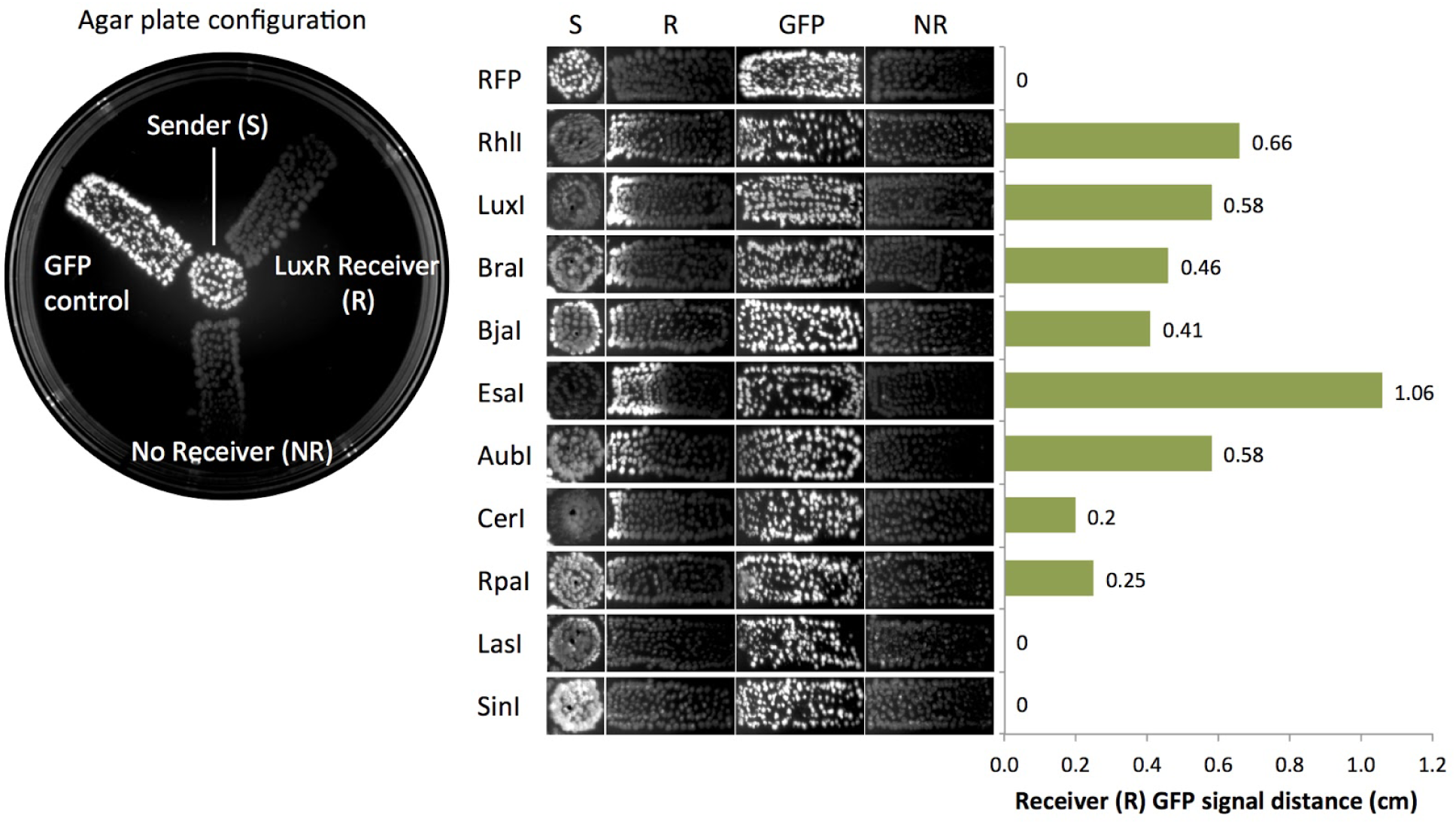
LuxR induction in solid agar cultures. The leftmost image shows the layout used to determine induction distance across agar. Sender bacteria were spotted in the center. Colonies transformed with Control-EGFP (GFP), receiver negative (NR), or LuxR receiver (R) plasmids were plated in streaks away from the center. (b) Induction distance in cm is shown for each of the ten senders tested.

**Figure 5:**
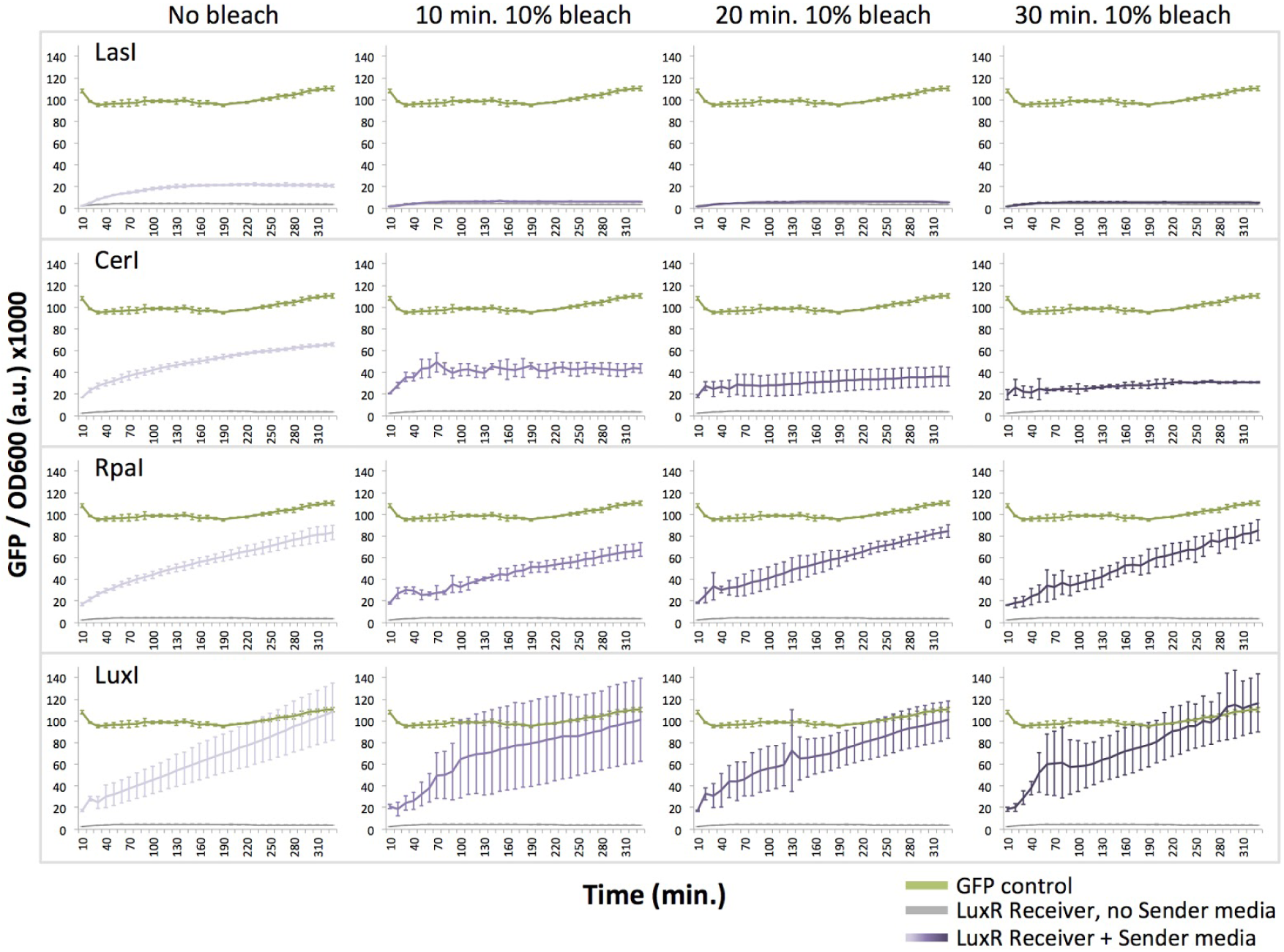
Bleach is not sufficient to deactivate media containing homoserine lactones (HSLs). Induction of the LuxR receiver device with untreated or treated media from LasI, CerI, RpaI, and LuxI sender cultures. Treated cultures were incubated for 10 min, 20 min or 30 min at a final concentration of 10% bleach.

### Comparison of conventional disposal methods: autoclaving is more effective than bleach for quenching HSL activity

We next tested whether standard hazardous waste treatment methods, bleaching and autoclaving, are sufficient to inactivate media containing HSLs from four Senders, LuxI, LasI, RpaI, and CerI [40]. These Senders span the range of LuxR-activation strengths we observed in the liquid culture experiments. Borchardt et al. showed that 3-oxo-containing HSLs are sensitive to oxidation in bleach solution, while other HSLs are resistant [41]. Therefore, we expected bleach to specifically quench the activity of HSL-enriched media from LuxI (3-oxo-C6-HSL) and LasI (3-oxo-C12-HSL).

We added bleach to a final concentration of 10% to overnight cultures of Sender-transformed *E. coli* BL21 for 10, 20, or 30 minutes. HSLs were extracted, added to liquid cultures of LuxR, and EGFP production was measured over time (330 minutes, or 5.5 hours). Consistent with the findings from Borchardt et al., we observed that bleach treatment eliminated the activity of the LasI-HSL-enriched media, while while media from CerI and RpaI remained active (Fig 5). Intriguingly, LuxI-HSL-enriched media showed no reduction in activity, which does not agree with the idea that oxidation destroys 3-oxo-C6-HSL activity. This result suggests the presence of bleach-resistant secondary HSLs or that the sterilization conditions used here are insufficient to destroy the activity of high concentrations of 3-oxo-C6-HSL. Overall, we conclude that treatment with 10% bleach for up to 30 minutes at room temperature is not a sufficient approach for quenching all HSL bioactivity.

To determine whether autoclave treatment is effective for deactivating HSLs, overnight cultures of *E. coli* BL21 transformed with the four sender plasmids were autoclaved for 15 min at 121˚C, 15 psi. HSLs were extracted and added to liquid cultures of LuxR immediately after autoclaving or after a 24 hour incubation at room temperature to represent the typical practice of storage between autoclaving and final disposal. For both immediate and delayed induction, autoclave treatment sufficiently inactivated HSLs produced by RpaI, CerI, and LasI senders (Fig 6). Autoclaved LuxI media showed some induction above background, however it is greatly reduced compared to untreated LuxI media. We conclude that autoclaving is the more effective treatment to deactivate media containing HSLs prior to disposal.

**Figure 6:**
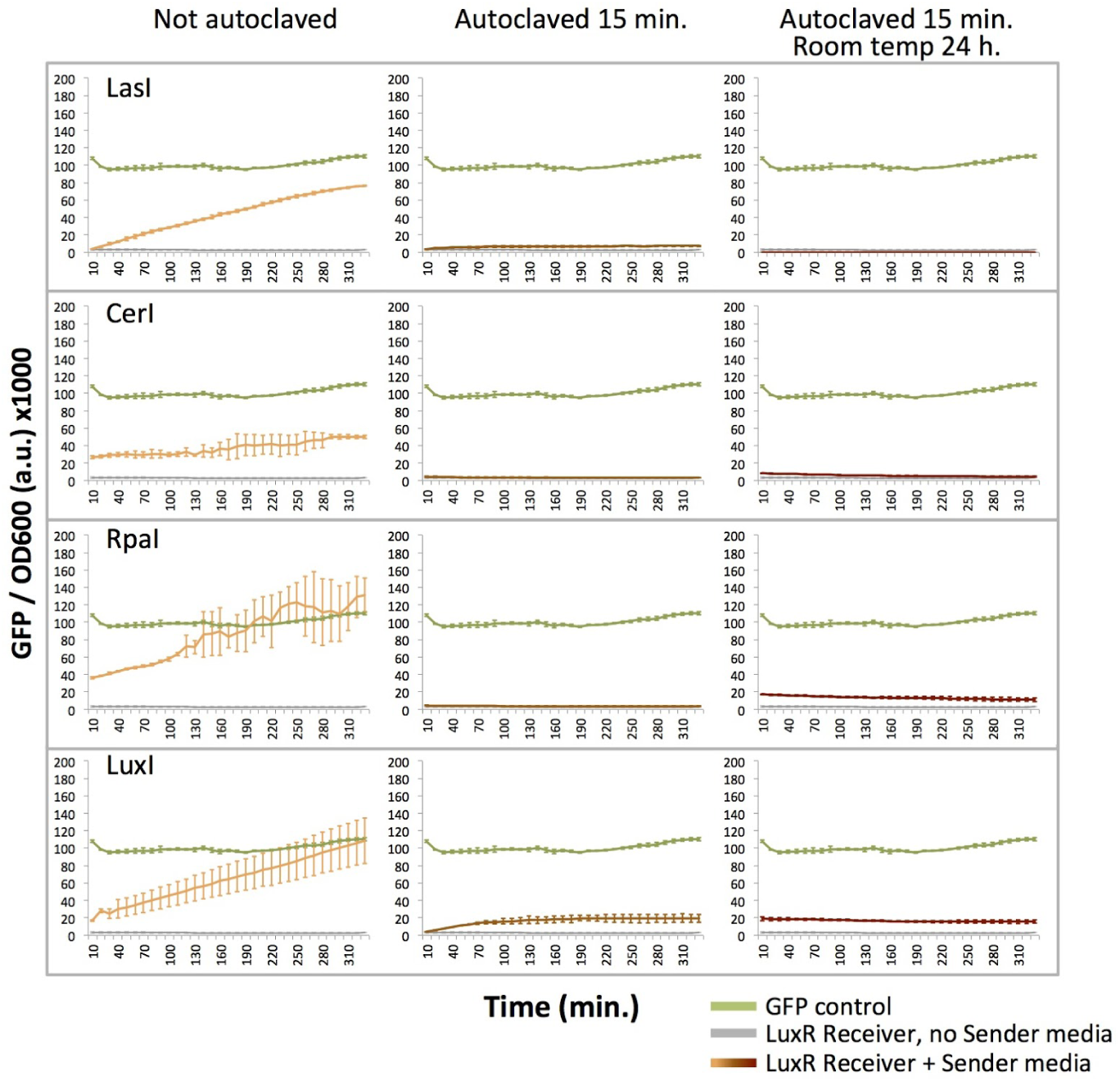
Autoclaving deactivates media containing homoserine lactones (HSLs). Induction of the LuxR receiver device with untreated or treated media from LasI, CerI, RpaI, and LuxI sender cultures. Treated cultures were autoclaved for 15 min. and used to induce LuxR receiver cultures immediately or after a 24 h incubation at room temperature.

## DISCUSSION

This study expands the quorum sensing toolbox by characterizing a library of ten HSL Sender devices in a commonly used lab chassis, *E. coli* BL21 in two contexts, liquid media and solid agar cultures. Using a Receiver device, LuxR, we confirmed functionality for nine of ten senders. By testing first in the chassis (*E. coli*) of interest at multiple dilutions, one can quickly gather data on how devices will behave in an *in vivo* circuit. Previous studies have demonstrated the orthogonality of LasI/LuxR in *E. coli* and RpaI/LuxR in *Salmonella typhimurium* and implemented circuits without crosstalk [8,13]. In our study, LasI and RpaI did not induce full activation (compared to Control-GFP) at low concentrations, and approached full activation (∼80%) at the highest concentration of HSL-enriched media. Assuming that Synthase expression can be tuned by using promoters and RBSs of different strengths, or origins of replication with different copy numbers, an engineer could exploit the varying sensitivity of a Receiver (e.g.LuxR) to mitigate crosstalk between parallel quorum sensing pathways.

Furthermore, synthases are known to produce different HSL profiles when expressed in non-native bacteria [31,39]. An HSL profile determined in one experiment (e.g., via mass spectroscopy or HPLC) may not be sufficient to predict the impact of a transgenic synthase on a Receiver. In our study, we selected synthases based on characterized HSL profiles, but determined their bioactivity using live cultures. Our approach does not require mass spectrometry, which can be expensive and inaccessible to certain researchers such as small labs, teaching labs, and iGEM teams.

Our work also demonstrates the importance of testing quorum sensing networks in the context of interest. Liquid culture induction results did not predict relative induction distances solid agar cultures. We observed a wide range of induction distances for synthases that induced similarly in liquid cultures and LasI, which induced LuxR at high dilutions in liquid culture, did not induce LuxR on the agar plate. These data confirm the need to test networks in context and provide researchers with data to inform design of agar-based devices using sender and receiver pairs that show high levels of crosstalk in other contexts. Interestingly, we did not see a gradient of induction on the agar plate inductions; for each sender tested, there was a defined edge where induction stopped. This is consistent with the behavior of the edge detection device built by Tabor *et al.* and suggests other senders could be used in this kind of device to create edges of varying thicknesses [42].

Finally, our work underscores the need for more careful consideration of disposal practices of HSL-enriched media. Risks associated with contamination by live cells are often considered and mitigated by sterilization and genetic containment [43]. In 2001, Borchardt et al. reported that acetylated homoserine lactones that lack the 3-oxo group failed to react with oxidized halogens (e.g., sodium hypochlorite, or bleach) and retained their ability to stimulate a QS-regulated gene in live cells while HSLs containing the 3-oxo group were effectively deactivated [41]. Consistent with their results, we found that cultures treated with bleach were still able to induce the LuxR receiver. We discovered that autoclaving sufficiently reduced bioactivity for all HSL-enriched media tested in our study. Further work should analyze a wider range of HSL to identify a universal sterilization strategy. We recommend that scientists who regularly use HSL-producing cultures perform their own bioactivity tests or autoclave their cultures before disposal. It is important to note that while we observed induction with bleached cultures in the lab, this does not suggest that environmental bacteria are being induced by disposed media. Our culture volumes were on the order of 1 mL to 100 mL which are immediately diluted by many orders of magnitude upon disposal. We have not provided evidence that additional safety precautions are required for using quorum sensing networks at small volumes and expressed in lab safe bacterial strains. Gene synthesis companies do not consider quorum sensing genes as hazardous when they are derived from a non-pathogenic bacteria. Many quorum sensing synthases are available in the iGEM registry and other repositories. We recommend that these organizations consider providing additional information to researchers on proper disposal.

In conclusion, our work adds to the rich, growing body of knowledge to inform design principles and best practices for the use of HSL-modulated quorum sensing networks in engineered circuits [18,19]. A series of recent publications has greatly expanded the number of HSL Receiver-type devices available to genetic engineers [18,30,44,45]. Our work to build and characterize a library of diverse senders complements these efforts to enable researchers to build more complex networks with minimal crosstalk, and provides an approach to identify functionally orthogonal sender-receiver pairs for parallel computation in multiple contexts.

## METHODS

### Plasmid Constructs

DNA digests included 1 μL (each) FastDigest enzymes and 1x universal buffer from Thermo Fisher Scientific and ∼2 - 4 μg DNA in a final volume of 30 μL, incubated at the lowest appropriate temperature for 10 min. Ligations included a 2:1 molar ratio of insert:vector, 1 μL T4 DNA ligase (New England Biolabs #M0202), and 1x Rapid ligation buffer (Roche #11635379001) in a final volume of 10 - 15 μL incubated at room temperature for 5 min. ***Inducible LuxR Receiver***: BBa_F2620 (iGEM Headquarters) contains the promoter pTetR controlling expression of the LuxR regulator and the LuxR-HSL regulated promoter pLuxR. A *XbaI-PstI* fragment (RBS-EGFP-2xTerminator) from BBa_E0240 was ligated downstream of pLuxR into a SpeI-PstI-linearized BBa_F2620 vector. ***Modular Sender Vector***: pTetR (BBa_R0040) was inserted upstream of an RBS (BBa_B0034) in pSB1A3 using BioBrick assembly [46]. pTetR-RBS was amplified with High Fidelity Phusion PCR (Thermo Fisher Scientific #F530, manufacturer‘s instructions) using primers 5‘-ctaggaattta<u>gtcttc</u>**tccctatcagtgata** (forward) and 5‘-<u>actagt</u>c[tctaga]agcggccgc[gaattc]**tttctcctctttctc** (reverse), then double digested with BbsI (to generate an *EcoRI*-compatible overhang 5‘-aatt) and SpeI. Similarly, a RBS-mCherry-2xTerminator fragment was amplified from BBa_J06702 (iGEM Headquarters) using primers 5‘-<u>actagt</u>**aaagaggagaaatac** (forward) and 5‘-ctag<u>ctgcag</u>a**tataaacgcagaaag** (reverse), then double digested with SpeI and PstI. Both fragments were ligated into a EcoRI-PstI-linearized pSB1A3 vector. For the preceding primer sequences: underlined text, restriction sites for cloning; bold text, template binding sequence; brackets, *EcoRI* and *XbaI* sites used for synthase ORF inserts (described below). Complete, annotated sequences for all Sender plasmids and the Receiver plasmid are available at the Haynes Lab Benchling website: https://benchling.com/hayneslab/f_/UWzcp3nk-quorum-sensing-collection

### Cloning of HSL Synthase Homologues

The coding regions for the following HSL synthases were synthesized as double-stranded oligos (IDT) with an *EcoRI* binding site upstream and a *XbaI* cut site downstream: AubI, BjaI, BraI, CerI, EsaI, LasI, LuxI, RhlI, RpaI, and SinI. The Modular Sender Vector and each HSL synthase oligo were cut with EcoRI and XbaI (Thermo Fisher Scientific) and ligated using T4 ligase (New England Biolabs). Plasmids containing synthases not already in the iGEM registry were submitted with the following identification keys: AubI BBa_K2033000, BjaI BBa_K2033002, BraI BBa_K2033004, CerI BBa_K2033006, SinI BBa_K2033008.

### Sender Media Preparation

Cells transformed with Sender plasmids or an mCherry (no synthase) plasmid were grown as colonies, and a single colony from each was used to inoculate 3 ml of LB with 5μg/ml ampicillin (VWR). Cultures were grown for 16 h at 37˚C with shaking. Sender and mCherry cultures were spun at 4500g for 5 min. Supernatants were passed through 0.22 μm nylon filters (VWR International) to remove any remaining cells.

### Microwell Plate Reader Assays

Single colonies of *E. coli* BL21 cells transformed with Receiver (F2620-EGFP) or Control-EGFP plasmid (pTrc99A vector expressing EGFP from the pLac promoter) were inoculated in 3 mL of LB broth [25g Acros LB broth Lennox granules (Sigma) in 1000 mL water] with 5 μg/mL ampicillin (VWR) and grown for 16 h at 37˚C with shaking. Each of culture was used to inoculate a new culture with a starting absorbance (OD600) of 0.05, and grown at 37˚C with shaking to a final OD600 of 0.8. Cultures were spun at 4500 xg for 5 min. Supernatants were discarded and cells were resuspended in fresh LB with ampicillin (OD600 = 0.8). Corning Black Costar Clear Bottom 96 Well Plates (Fisher Scientific) were loaded with 150 μL Receiver or Control-EGFP of a total volume of 300 μL per well with a final OD600 of 0.4. Varying concentrations of sender and negative sender media were used to ensure the same total volume of spent media per well (Table 1). In addition to a total of 150μl spent media, 150 μL of working receiver stock was added per well. Plates were analyzed on a Biotek Synergy H1 Microplate Reader, measuring red fluorescence (580 nm to 610 nm), green fluorescence (485 nm to 515 nm), and OD600 every 10 min for 8h at 37˚C with shaking. Automatic gain adjustment was set to scale to the lowest detected well values for each measurement. Mean values and standard deviations of GFP signal / OD600 (a.u.) were calculated for triplicate wells at each 10 min time point. Graphs and heat maps were generated in Microsoft Excel 2016.

### Agar Plate Inductions

Cultures of *E. coli* BL21 (New England Biolabs) transformed with Sender plasmids, a Receiver plasmid, a negative receiver plasmid, and a GFP positive plasmid were grown in 3 mL of LB with 5 ug/mL ampicillin for 16 hours at 37˚C and shaking at 220 rpm. Bacterial culture was subsequently spread onto LB agar supplemented with 5 ug/mL ampicillin with sterile disposable plastic micropipette tip, such that a central spot of sender culture would evenly diffuse towards proximal receiver and Control-EGFP positive control cultures. Plates were grown for 16 hours at 37˚C. Images were acquired with a Pxi4 imager under ultraviolet light, saved at 300 dpi resolution, and analyzed using ImageJ software. Edges of fluorescence-positive areas were determined as the area in which the raw integrated density within a window of 50×100 pixels was equal to that of GFP-minus cells. Induction distances were determined as the shortest distance (straight line) from the Sender-proximal edge of the Receiver cells to the edge of the fluorescence positive area.

### Bleach and Autoclave Treatment of Sender Media

HSL-enriched media (prepared as described above) from all senders was treated with bleach (sodium hypochlorite solution) at a final concentration of 10%. Bleach treatment was performed by adding 1 volume of bleach (Genesee Scientific) to 10 volumes of filter-purified, HSL-enriched medium. Samples were incubated at room temperature for 10, 20 or 30 min. At each time point, 1 mL of solution was removed and added to an equal volume of ethyl acetate for HSLs extraction. The extraction solution was incubated for 1 minute at room temperature while shaking, and then allowed to phase separate for 10 min at room temperature. The organic phase was collected and subjected to rotary evaporation. The dried samples were resuspended with 1 mL of sterile, double distilled H2O. A control sample of sender media without bleach was also extracted at each time point. For autoclave treatments, 1 mL of the HSL-enriched media was sealed in a glass round-bottom tube and autoclaved for 15 min at 121˚C and 15 psi (BetaStar 26 x 26 x 39 Automatic Vertical Sliding Autoclave). After cooling, samples were extracted as described above. Receiver inductions were carried out on treated and extracted HSL-enriched media as described above.

## Author Contributions

RD and CB designed and optimized the Receiver induction strategy. RD designed, built, and verified the plasmid constructs with assistance from RM and JW. EL, JW, JX, BD, and RM performed preliminary plate reader assays. ST generated the final data shown in Fig 3. EL performed the agar plate assays. EL, JW, JX, and BD completed the bleached and autoclaved media experiments with supervision from RD. RD, CB, and ST supervised work done by the undergraduate students. KAH oversaw the completion of the work, created the final figures, and completed final editing of the manuscript. All authors approved the final version of the manuscript.

## Acknowledgements

Most of the work reported here was presented at the 2016 International Genetically Engineered Machines (iGEM) Competition by the Arizona State University (ASU) 2016 iGEM team. The authors thank J. Steel (DNASU) for DNA sequencing services. The authors thank R. Allen, S. Raman, G. Srinivasan and J. Daer for critical review of the manuscript. The authors thank Integrated DNA Technologies (IDT), iGEM Headquarters, the Western Alliance to Expand Student Opportunities (WAESO), the School of Molecular Sciences at Arizona State University, and the School of Biological and Health Systems Engineering at Arizona State University for their support of the undergraduate co-authors. KAH was supported by the NIH NCI (K01CA188164). RMD was supported by the ARCS Foundation and NSF CBET (1404084). CMB is supported by the Ira A. Fulton School of Engineering at Arizona State University.

